# Could Italy host the new coronavirus before China?

**DOI:** 10.1101/2021.02.24.432649

**Authors:** Aleš Tichopád, Ladislav Pecen

## Abstract

The pandemic of the COVID-19 disease caused by the SARS-CoV-2 virus has been believed to originate in China and spread later to other parts of the world. It is well acknowledged that the first diseased individuals appeared in China as early as in December 2019, and possibly even earlier in November. It has also been well established that the virus stroke Italy later in January or in February 2020, distinctly after the outbreak in China. Paper by Apolone et al. published in a local Italian medical journal in November 2020 however exposed this chronology to doubt.

By fitting early part of the epidemic curve with the exponential model and extrapolating it backwards in time, we could estimate the day-zero of the epidemic and calculated its confidence intervals in Italy and China. We also calculated how probable it is that Italy encountered the virus prior 1 January 2020.

We determined an early portion of the epidemic curve representing unhindered exponential growth which fit the exponential model with high determination >0.97 in both countries. We suggest that the day-zero in China and Italy was 8 December (95% CI: 3 Dec., 20 Dec.) and 22 January (95% CI: 16 Jan., 29 Jan.), respectively. We could with high confidence reject that Italy encountered the virus earlier than China (p <0.01).

Based on our analysis we oppose the findings published by Apolone at al. and view the proposed pre-pandemic presence of the virus in Italy as very unlikely.

## Introduction

On 14 January, after months of negotiations between the WHO and Beijing, a WHO team arrived in China with the goal to set up a research probing origin of the SARS-CoV-2. This effort bearing significant political and economic dimension underlines the persisting dispute over the origin of the virus and the responsibility for its uncontained global spread. On February 9 the mission was completed with no break-through findings.

The global pandemic of the COVID-19 disease caused by the SARS-CoV-2 virus will certainly be tracked among the most significant global events in the recent history with a death toll nearing 2.5 Million as of the end of January 2021 [1]. Its humanistic, economic, and political impact has not only shaped the ongoing efforts to contain its spread but also relationships between countries, their citizens, and politicians.

According to a study by Huang et al. [2] the first case of COVID-19 dates to December 1, 2019 but other sources propose there may have been patients exhibiting same symptoms already in November of the same year [3,4]. Reported by the South China Morning Post, the first person with confirmed COVID-19 dating back to 17 November 2019 was a 55-year-old male patient from the province of Hubei [5]. This report further said Chinese authorities had by the end of the year identified at least 266 people who contracted the virus and who came under medical surveillance. Interestingly, none of these first reported patients have direct link with the Wuhan Seafood Market that has been associated with the origin of the virus as late December Chinese doctors came to realize that they were dealing with a new and serious virus in increasing number of patients with similar symptoms mostly originating from Wuhan [6]. On the early December days Chinese hospitals reported one to five cases with similar symptoms each day. According to information conveyed to WHO by Chinese authorities on 11 and 12 January, 41 cases with novel coronavirus infection have been preliminarily diagnosed in Wuhan. Symptom onset of the 41 confirmed COVID-19 cases ranges from 8 December 2019 to 2 January 2020. No additional cases have been detected from 3 to 12 January 2020 [6]. Further on 11 January, the first SARS-CoV-2 virus genome sequence was deposited in the GENBNK (the NIH database with public genetic sequences) and shared with public via virologist.org and uploaded to the platform GISAID [7]. On the following day, the information on discovery of the genome sequence was officially shared with the WHO. In parallel, on 12 January Chinese Health authority closed the laboratory that was the first to share the coronavirus genome with the world. On 23 January, the first coronavirus lockdown came into force in Wuhan. Mass testing, mask use enforcement and varying degree of social distancing measures helped to gain an effective control over the epidemics within few weeks in China.

Italy became the first epicentre of the COVID-19 outside China. The outbreak in Italy officially started in Rome on 31 January, after two Chinese visitors who arrived from Wuhan on 23 January, tested positive for SARS-CoV-2 [8]. They later recovered and no further person was identified infected through their contacts. Later however, following admitted failures in handling a COVID-19 patient by hospital in the northern town of Codogno, Lombardy, 322 cases were reported by the Italian civil protection agency on 25 February. On 21 February, a 38-year-old man was admitted to the hospital in Codogno and was confirmed as the first Italian citizen with COVID-19 reported later as the patient 1. In few days, the number of COVID-19 cases increased beyond expectations. Italy became, to that date, the country with the highest number of officially reported and confirmed COVID-19 patients outside Asia. The disease initially swept through Lombardy, a northern region of Italy, and spread in a lesser extent over the whole country. A lockdown was triggered on 8 March in the most affected north of the country, and extended gradually until 4 May, the date on which relaxation of the measures commenced. The massive outbreak in Lombardy put the regional health service under considerable strain. There has been no evidence suggesting Italy could encounter the virus already in 2019. The only available published analysis to establish the day-zero in Italy suggests it was 14 January [9].

Throughout the first half of 2020, facing the threat of a new unknown disease and the adverse impact of containment measures, a critical public view has formed directed against those considered responsible for the initial loss of control over the spread. China has come into the light of worldwide public criticism. Stung by complaints it allowed the disease to spread, Chinese political representatives claimed the virus came from abroad and it is therefore not the “China virus”. Along with the rapid global spread of the pandemic later in 2020 and with the onset of the autumn waves in Europe, the European public frustration gradually transformed into different forms, including denialism, defiance and anti-vaccination movement. After the initial shock in the first two months of 2020, disinformation and scientifically unfounded theories of all kind started soon proliferating in social as well as mainstream media, present various views of the source and the origin of the virus including the lab release theory. These narratives were to a great degree amplified by the US official sources including US embassy in China, the former US President and the Office of the director of national intelligence (ODNI). ODNI on 30 April specified that US intelligence officials were investigating whether the outbreak began through contact with animals or through a laboratory accident. Approximately from May on, also less sophisticated disinformation filled internet and some media, proposing a man-made theory pointing, for instance, at the US military as the engineer of the virus. Many other more or less bizarre hoaxes on the origin of the virus followed and have persisted until today [10].

An unexpected detection of SARS-CoV-2 antibodies in the pre-pandemic period was reported by Italian research team in a rather low impact factor journal in November 2020, dating some positive tissue samples back to September 2019 [11]. This information shed light on the possibility that the virus had been spreading in Italy well before the outbreak was officially reported in Wuhan, China. However, this information did not seem to dramatically shake the existing view of the virus origin, it soon populated media worldwide. Most of the expressed scientists’ scepticism pointed on the specificity of the antibody tests used. No peer-reviewed paper addressing this or other aspects of the paper by Apolone et al. [11] has been however published to date.

We want to respond to speculation that the virus could be in Italy before the outbreak in China, which appeared as a result of the above-mentioned publication by Apolone et al. [11].

## Methods

### Log-linear analysis

The data on new daily cases were taken from the web of the European Centre for Disease Prevention and Control [12]. We identified the earlies consistent portion of the epidemic curve in both China and Italy, which we believe describes the best the epidemic progression within a naive population with minimum imposed measures affecting its intrinsic dynamics. It was repeatedly shown that epidemic curve in its early portion can be well fitted by simple phenomenological models as intrinsic growth is the determining variable [13,14]. Considering the interval from January through April only, we identified the most linear early portion of new cases projected day-by-day on log-linear plot, using a semiautomatic process. In SAS v. 9.4 for MS Windows, we stepwise rolled a 10-day window over the data and determined the root mean square error (RMSE) from a linear regression fitted within each window. Eventually, the 10-day interval with the smallest RMSE was considered the most linear and was subsequently analysed by extending the 10-day window on both ends by adding one, two or three additional data points and recalculating the RMSE. This process was repeated if it diminished the RMSE obtained or maintained it constant at least.

### Model fitting

Using proc *nlin* in SAS we fitted a simple exponential model to the above-described delimited data on a linear/linear scale

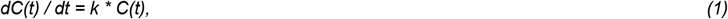

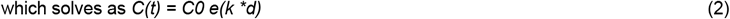

where *C* denotes the daily number of new cases on day *d*, and *k* determines the growth. This is a fundamental model to describe the early ascending progression of the epidemic due to its intrinsic dynamic in an environment reach in naive host. On a semi-logarithmic scale, exponential growth pattern is visually evident if a straight line fits well several consecutive days. The model 1 is an established elementary principle in epidemiology [13,14].

The proc *nlin* of the SAS program fits nonlinear regression models and estimates by means of iterative process (here the Marquardt method was chosen) the parameters by nonlinear least squares. The determination of each fit was obtained analogically to linear regression as 1 – variance of model residuals / variance of the y variable (here the reported new cases C).The proc *nlin* also produces predicted values for missing entries prior, within, and after the fitted area.

### Location of day-zero

Having the model with all parameters determined, the day-zero (*d*_*0*_) is located where *C(d)* rounded to the nearest whole number equals zero on a given day while it equals one on the next day. To determine the probability of the *d*_*0*_ being prior a specified day, the alpha level of the fitted model equation was automatically iterated to establish the lower confidence interval alpha at the specified day, here 1 January 2020. Then the probability is given as 2-times alpha since only one-sided probability is associated with the question whether the Italian day-zero could precede the 1 January.

In addition, we looked at 95% and 99% confidence interval for the d_0_ estimates in both countries to judge the probability that Italy hosted the virus before China.

## Results

Performing the log-linear analysis on the Italian data, we identified the interval from 28 February through 15 March as the early conservative undeflected growth portion of the epidemic curve, comprising 17 days. Similarly, for China we identified the interval from 29 January through 6 February, in total only 9 days (Figure 1). However, for the lack of data points, we decided to retain also all the early data points prior 29 January to be fitted with the primary model for China (full data, N=24).

**Fig 1.**
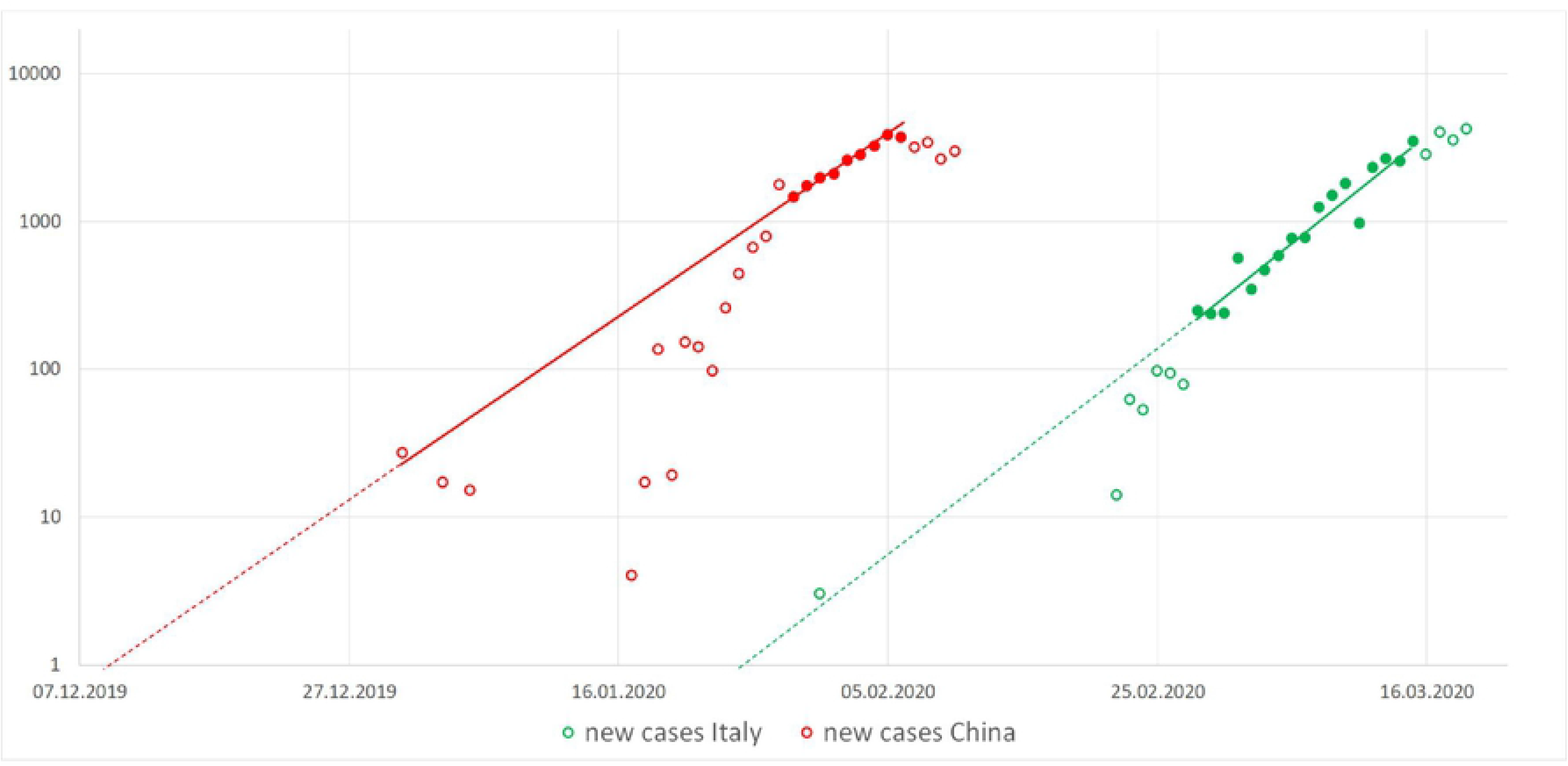
Log-liner plot of new cases with the linear period represented by the dots identified by the semiautomatic process. Data represented by the full dots could be identified as exponentially behaved.

Using the simple exponential model 1 we estimate that the *d*_*0*_ in China and Italy was 8 December 2019 (95% CI: 3 Dec., 20 Dec.) and 22 January (95% CI: 16 Jan., 29 Jan.), respectively. The model could be well fitted over both the Chinese as Italian data with high determination R^2^ equaling 0.973 and 0.975, respectively (Figure 2).

**Fig 2.**
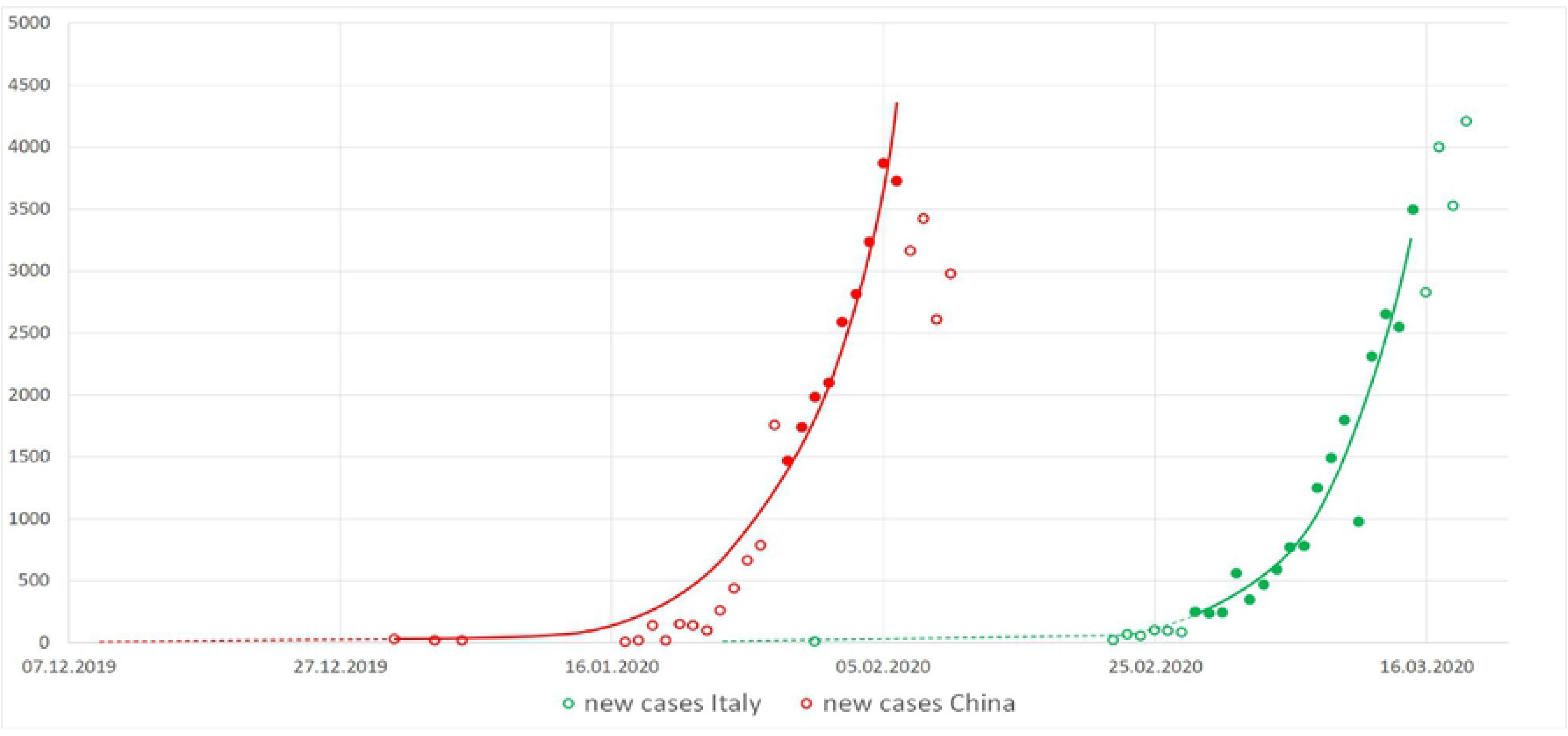
Exponential fit (full line) over a selected portion of data. Data represented by the dots could be identified as exponentially behaved.

As a sensitivity analysis we trimmed the early data from the Chinese curve which departs from the log-liner trend analysis. Trimming off the early points from the full data and fitting the remaining N=9 data moved the estimated *d*_*0*_ 11 days ahead to 27 November. The model could be fitted with even higher determination 0.997 than the full data model, however with greater uncertainty of the estimated *d*_*0*_ (95% CI: 31 Oct., 22 Dec.).

To address our objective – comparisons of *d*_*0*_ between China and Italy– we conservatively consider the 8 December 2019 as the *d*_*0*_ in China as produced by the full data fit. The 99% confidence interval for the estimated *d*_*0*_ in both China and Italy were obtained as (99% CI_China_: 29 Nov., 24 Dec.) and (99% CI_Italy_: 11 Jan. 2020, 3 Feb. 2020), respectively. The 99% confidence intervals do not overlap, showing hence two distinct and separate onsets of the epidemics, whereas the first occurred in China. This conclusion remains unchanged also when the *d*_*0*_ based on the trimmed Chinese data with N=9 is considered (99% CI: 16 Oct., 26 Dec.)

Based on the model estimates the probability that Italy encountered the infection already prior 1 January 2020 is calculated as low as 0.00004 (0.004%).

## Discussion

The virus showed that it can only be combated successfully if well informed strategies are proposed by expert teams and adopted by countries and broad regions such as the EU. Erroneous or misleading information can bias scientific, political or civic opinions and jeopardise proposed strategies to combat the pandemic. Given its magnitude, it is almost certain that human lives will be paid as the ultimate toll for inadequate policy. As many papers regarding the global pandemic are published under some kind of a fast-track process, critical review of published information is critical and should be taken up by broader scientific community. Deep understanding of a pre-pandemic virus pathways is of great importance for better preparedness and response to future outbreaks.

We made an attempt to revise the speculation that Italy could host SARS-CoV-2 prior China has confirmed the outbreak. However may this seem to be built on a weak foundation to anybody who remembers news headlines from the turn of 2019 and 2020, all pointing at China, it deserves attention in the light of recent findings [11]. To address the persisting uncertainty nourished by various politically motivated narratives and media, WHO launched an international investigation to assemble evidence on the geographical and biological origin of the virus and its pre-pandemic trajectory [15].

Given our aim was a backwards extrapolation of the earliest day when a local inhabitant encountered the virus, a simple exponential model proved suitable to characterize the early phase of the epidemic progressing without hindrance. We could sufficiently well identify a consistent exponential growth period from of the epidemic curve in Italy using a semiautomatic process on a log-linear scale. In Italy, this period spanning from 28 February through 15 March can be associated with the imposed lockdown in Lombardy which started on 8 March, assuming the lockdown triggered an effect that shaped the data few days later. Less tight association we can see between the detected early exponential period 29 January through 6 February, which is distinctly later than the onset of the lockdown in China on 23 January. We explain this by the generally higher uncertainty of the data from China.

Our localization of the day-zero of the Italian outbreak supports the existing chronology of the early outbreak in Italy with the first significant cluster of patients reported in the second half of February [8,9]. The work by Russo et al. suggests 14 January as the day-zero in Italy, which is eight days earlier than our estimate and within our 99% confidence interval (11 January, 3 February). The difference could be explained by the delay between the onset of the disease and the date of diagnostics. In our model we did not adjust our estimated day-zero for this delay as, for the early period of the outbreak, we found no reliable data to support it. The Italian outbreak was initially concentrated in the north of the country and hence the data we used for fitting come predominantly from this regional outbreak and does not compound several parallel outbreaks. It was first later in March and April when the epidemics spread over the entire country and abroad [16]. This further justifies the use of a simple model over a compounded one. The breakout in the north of the country was believed to be likely initiated by a single supper-spreader while at that time there was no case detected elsewhere in Italy. The dynamic of the subsequent epidemics within a naive population shows that the virus could quickly exploit the new ecological opportunity and it is very unlikely, if not merely impossible, that any such previous opportunity would have been missed in the past unless there were a stringent containment measures in place.

In the paper by Apolone at al. [11], among 959 volunteers participating on a prospective lung cancer screening trial between September 2019 and March 2020 in Italy positive serologic assay probes have been reported in 111. Several tens of the positive tested individuals were reported as early as in September and October 2019 and seem to have been located across the whole country, yet with some degree of overlap with the incidence spatial pattern of the later COVID-19 outbreak in 2020. None of those 111 individuals was diagnosed with COVID-19 nor they reported symptoms. They were presumably living under conditions facilitating regular social contact. Not combated by any means of social distancing at that time, presence of SARS-CoV-2 virus among interacting individuals in September would inevitably lead to a devastating nation-wide outbreak earlier than the in March, most likely already on December with a notable excess of all-cause mortality far exceeding its maximum reported in week 14 of 2020 [17]. Nothing like that has been observed though. It is also strikingly high prevalence (111 out of 959 means 11,6%) reported in the cohort studied by Apolone at al. compared to 2.6% reported in February 2020 among 2,812 individuals [11, 18].

Much has been written and said about specificity of various SARS-CoV-2 antibody-based assays and it is out of scope of this paper and competence of its authors to deep dive into this topic. One of several possible explanations of the surprising finding by Apolone et al. could consist in the use of a cross-reaction of the used antibodies [19].

The strength of our study is its robustness consisting in the use of adequate model on a consistent data with minimum assumptions needed. The main limitations of our study is the unknown proportion of undetected infected individuals and the unknown time between the infection and the test date in the early days of the pandemic. Assuming that both would have a substantial effect, it would shift the day-zero by days. We believe that this error would rather constantly affect both countries and hence more or less preserve the order.

In our work no medical or research intervention was imposed on humans or animals. We did not specifically address individual human subjects.

## Authors’ contributions

AT: Preparation of the overall concept and the methodical approach, literature review and writing, review of the statistical analysis. LP: Statistical analysis, statistical methods writing and critical review of the manuscript.

## References

1. Institute for Health Metrics and Evaluation (IHME). COVID-19 Projections. Seattle, WA: IHME, University of Washington, 2020. [Accessed: 1 Feb 2021]. Available from https://covid19.healthdata.org/projections (28 January 2021).

2. Huang C, Wang Y, Li X, Ren L, Zhao J, Hu Y, et al. Clinical features of patients infected with 2019 novel coronavirus in Wuhan, China. Lancet. 2020;395(10223):497–506.

3. Bryner J. 1st Known Case of Coronavirus Traced Back to November in China. 2020; [Accessed: 25 Jan 2021]. Available from: https://www.livescience.com/first-case-coronavirus-found.html.

4. Davidson H. First Covid-19 Case Happened in November, China Government Record Show – report. 2020; [Accessed: 25 Jan 2021]. Available from: https://www.theguardian.com/world/2020/mar/13/first-covid-19-case-happened-in-november-china-government-records-show-report.

5. Ma J. Coronavirus: China’s First Confirmed Covid-19 Case Traced Back to November 17. 2020; [Accessed: 25 Jan 2021]. Available from: https://www.scmp.com/news/china/society/article/3074991/coronavirus-chinas-first-confirmed-covid-19-case-traced-back.

6. WHO. Emergencies Preparedness, Response: Novel Coronavirus – China. 2020; [Accessed: 28 Jan 2021]. Available from: https://www.who.int/csr/don/12-january-2020-novel-coronavirus-china/en/.

7. ECDC. Timeline of ECDC’s reponse to COVID-19. 2020. [Accessed: 25 Jan 2021]. Available from: https://www.ecdc.europa.eu/en/novel-coronavirus/event-background-2019.

8. Carinci F. Covid-19: preparedness, decentralisation, and the hunt for patient zero. BMJ 2020; 368.

9. Russo L, Anastassopoulou C, Tsakris A, Bifulco GN, Campana EF, Toraldo G, et al. Tracing day-zero and forecasting the COVID-19 outbreak in Lombardy, Italy: A compartmental modelling and numerical optimization approach. PLoS One. 2020;15(10).

10. Tagliabue F, Galassi L, Mariani P. The “Pandemic” of Disinformation in COVID-19. 2020 SN Compr Clin Med. Aug 1:1–3.

11. Apolone G, Montomoli E, Manenti A, Boeri M, Sabia F, Hyseni I, et al. Unexpected detection of SARS-CoV-2 antibodies in the prepandemic period in Italy. Tumori. 2020;

12. ECDC. Download historical data (to 14 December 2020) on the daily number of new reported COVID-19 cases and deaths worldwide. 2020. [Accessed: 27 Jan 2021]. Available from: https://www.ecdc.europa.eu/en/publications-data/download-todays-data-geographic-distribution-covid-19-cases-worldwide

13. Ma J. Estimating epidemic exponential growth rate and basic reproduction number. Infect Dis Model. 2020; 5:129–141.

14. Chowell G, Sattenspiel L, Bansal S, Viboud C. Mathematical models to characterize early epidemic growth: A review. Phys Life Rev. 2016;18:66–97.

15. Mallapaty S. Where did COVID come from? WHO investigation begins but faces challenges. Nature. 2020; 587(7834):341–342.

16. Gatto M, Bertuzzo E, Mari L, Miccoli S, Carraro L, Casagrandi R, Rinaldo A. Spread and dynamics of the COVID-19 epidemic in Italy: Effects of emergency containment measures. Proc Natl Acad Sci U S A. 2020;117(19):10484–10491.

17. EuroMOMO. Graphs and maps. 2021. [Accessed: 30 Jan 2021]. Available from: https://www.euromomo.eu/graphs-and-maps

18. Lavezzo E, Franchin E, Ciavarella C, Cuomo-Dannenburg G, Barzon L, Del Vecchio C, et al. Suppression of a SARS-CoV-2 outbreak in the Italian municipality of Vo’. Nature. 2020; 584(7821):425–429.

19. Vandenberg O, Martiny D, Rochas O, van Belkum A, Kozlakidis Z. Considerations for diagnostic COVID-19 tests. Nat Rev Microbiol. 2020:1–13.

